# Cortical processing of breathing perceptions in the athletic brain

**DOI:** 10.1101/174052

**Authors:** Olivia K. Faull, Pete J. Cox, Kyle T. S. Pattinson

## Abstract

Athletes regularly endure large increases in ventilation, and accompanying perceptions of breathlessness. While breathing perceptions often correlate poorly with objective measures of lung function in both health and clinical populations, we have previously demonstrated closer matching between subjective breathlessness and changes in ventilation in endurance athletes, suggesting that athletes may be more accurate during respiratory interoception. To better understand the link between exercise and breathlessness, we sought to identify the mechanisms by which the brain processing of respiratory perception might be optimised in athletes.

Twenty endurance athletes and 20 sedentary controls underwent 7 Tesla functional magnetic resonance imaging. Inspiratory resistive loading induced conscious breathing perceptions (breathlessness), and a delay-conditioning paradigm was employed to evoke preceding periods of anticipation. Athletes demonstrated anticipatory brain activity that positively correlated with resulting breathing perceptions within key interoceptive areas, such as the thalamus, insula and primary sensorimotor cortices, which was negatively correlated in sedentary controls. Athletes also exhibited greater connectivity between interoceptive attention networks and primary sensorimotor cortex. These functional differences in athletic brains suggest that exercise may optimise processing of respiratory sensations. Future work may probe whether these brain mechanisms are harnessed when exercise is employed to treat breathlessness within chronic respiratory disease.

## Introduction

Athletes are able to undertake incredible feats of human achievement, with faster, higher and stronger performances recorded each year. Whilst exercise training is known to induce widespread physiological changes in the periphery, the concurrent changes in the structure and function of the athletic brain are less well investigated. For endurance athletes, exercise training is targeted to improve the ability of tissues to utilize oxygen in the combustion of fuels such as fat and carbohydrate, producing the energy required for repeated skeletal muscle contraction (Holloszy & Coyle, 1984; Jones & Carter, 2000). However, the role of the brain in perceiving and modulating changing sensations from the periphery, useful for maintenance of homeostasis during situations of perturbed physiology, is often overlooked.

Ventilation during exercise is tightly controlled, balancing neurally-modulated feed forward ventilatory commands and peripheral feedback to stimulate appropriate ventilation for exercising needs (Kaufman & Forster, 1996; Waldrop *et al.*, 2010). Interoceptive monitoring of respiratory sensations contributes to the maintenance of homeostasis (Davenport & Vovk, 2009), and with sufficient exercise intensity, the strain of immense increases in ventilation induces perceptions of breathlessness (El-Manshawi *et al.*, 1986; Takano *et al.*, 1997; Lansing *et al.*, 2000; Borg *et al.*, 2010). While endurance athletes are repeatedly exposed to these respiratory sensations and breathlessness, it is as yet unknown whether brain networks involved in these perceptions may also adapt to better cope with exercise demands. This understanding would allow us to explore how processing of ventilatory signals might be altered in different states, such as in athletes or conversely in chronic respiratory disease, where subjective reports of breathlessness are often discordant with objective measures of lung function and ventilation (Herigstad *et al.*, 2017).

Importantly, prior experiences of strong respiratory sensations may also alter the way someone anticipates and perceives their breathing (Faull *et al.*, 2017; Van den Bergh *et al.*, 2017; Herigstad *et al.*, 2017). Expectations regarding upcoming respiratory sensations from conditioned cues (Pavlov, 1927), for example the breathlessness associated with an approaching hill whilst running, can be an important influence on both threat behaviours and preventative actions (i.e. to avoid the hill) (Lang *et al.*, 2011), or on the perception itself (Price *et al.*, 1999; Porro *et al.*, 2002; Wager *et al.*, 2004). Repeated breathlessness exposure may alter this anticipation in athletes, focusing their attention towards respiratory sensations (Merikle & Joordens, 1997; Phelps *et al.*, 2006; Ling & Carrasco, 2006), reducing their anxiety (Spinhoven *et al.*, 1997; Bogaerts *et al.*, 2005; Tang & Gibson, 2005) or improving their interoceptive ability (Gray *et al.*, 2007; Critchley *et al.*, 2013; Mallorqui-Bague *et al.*, 2016; Garfinkel *et al.*, 2016*b*; 2016*a*). Interestingly, exercise therapy is currently the most effective treatment for breathlessness associated with chronic obstructive pulmonary disease (COPD), improving breathlessness intensity and anxiety (Carrieri-Kohlman *et al.*, 1996; 2001; Herigstad *et al.*, 2017), without concurrent improvements in lung function. It is possible that athletes may have different prior expectations and anticipation of breathlessness, although this has yet to be investigated.

In previous work we have observed closer matching between changes in ventilation and perceptions of breathlessness in endurance athletes compared to sedentary individuals (Faull *et al.*, 2016*a*). Here, we sought to identify how the brain processing of both anticipation and perception of respiratory sensations may be altered in these athletes, to better understand potential contributors to ventilatory interoception. We investigated functional brain activity using magnetic resonance imaging (fMRI) during both conditioned anticipation and perception of a breathlessness stimulus. We also examined potential differences in the resting temporal coherence, or ‘functional connectivity’ (Gerstein & Perkel, 1969; Van Den Heuvel & Pol, 2010) of brain networks involved in attention towards sensory information, allostasis and interoception (Kleckner *et al.*, 2017). Differences in underlying functional connectivity may help us to understand how the athlete brain may be altered to facilitate accurate respiratory perceptions, and we hypothesized that these athletes would demonstrate both altered functional breathlessness-related brain activity and connectivity compared to their sedentary counterparts.

## Materials and Methods

### Subjects

The Oxfordshire Clinical Research Ethics Committee approved the study and volunteers gave written, informed consent. Forty healthy, right-handed individuals undertook this study, with no history of smoking or any respiratory disease. This cohort comprised two groups; 20 subjects who regularly participated in endurance sport, and 20 age- and sex-matched (±2 years) sedentary subjects (in each group: 10 males, 10 females; mean age ± SEM, 26 ± 1.7 years). Athletes were active participants in endurance sports (cycling, rowing and endurance running), with training sessions conducted at least 5 times per week. Sedentary subjects did not partake in any regular exercise or sport. Prior to scanning, all subjects underwent breathlessness testing during exercise and chemostimulated hyperpnea, which have been presented elsewhere (Faull *et al.*, 2016*a*), and a combined whole-group analysis of fMRI data has been previously reported (Faull & Pattinson, 2017).

### Stimuli and tasks

Subjects were trained using an aversive delay-conditioning paradigm to associate simple shapes with an upcoming breathlessness (inspiratory resistance) stimulus (Faull & Pattinson, 2017). A breathing system was used to remotely administer periods of inspiratory resistive loading to induce breathlessness (as predicted by the conditioned cues). The breathing system contained an inspiratory resistance arm (using a porous glass disk) with a non-rebreathing valve connected to a mouth piece, which could be periodically applied using the addition or removal of medical air through an alternative inspiratory non-rebreathing arm (detailed in (Faull et al., 2016b; Faull & Pattinson, 2017)). Mean peak inspiratory resistance was recorded at 14.7 (±8.3) cmH_2_O for the loading periods across subjects, and group values are presented in Tables 1 and 2. The subject’s nose was blocked using foam earplugs and they were asked to breathe through their mouth for the duration of the experiment.

**Table 1.** Mean (±sd) physiological variables across conditioned respiratory tasks. *Significantly (*p* < 0.05) different from sedentary group. Abbreviations: P_ET_CO_2_, pressure of end-tidal carbon dioxide; P_ET_O_2_, pressure of end-tidal oxygen; RVT, respiratory volume per unit time; bpm, beats per minute.

|  | Unloaded breathing |  | Anticipation |  | Breathlessness |  |
| --- | --- | --- | --- | --- | --- | --- |
|  | ATHLETE | SEDENTARY | ATHLETE | SEDENTARY | ATHLETE | SEDENTARY |
| $P_{ET}CO_2$ (mmHg) | 35.96 (5.56) | 35.08 (3.20) | 35.50 (5.81) | 34.76 (3.60) | 36.34 (6.23) | 35.40 (3.92) |
| $P_{ET}O_2$ (mmHg) | 129.68 (6.41) | 134.09 (15.15) | 129.55 (6.75) | 133.59 (13.47) | 131.18 (6.83) | 137.55 (16.42) |
| Respiratory rate ($min^{-1}$) | 10.15 (2.59)* | 13.35 (3.51) | 9.99 (2.63)* | 12.93 (4.29) | 9.40 (3.58) | 11.54 (5.11) |
| RVT (% change from baseline) | -4.06 (5.70) | -0.56 (7.94) | -0.03 (12.14) | 6.07 (18.78) | -20.00 (24.88) | -13.23 (28.54) |

**Table 2.**
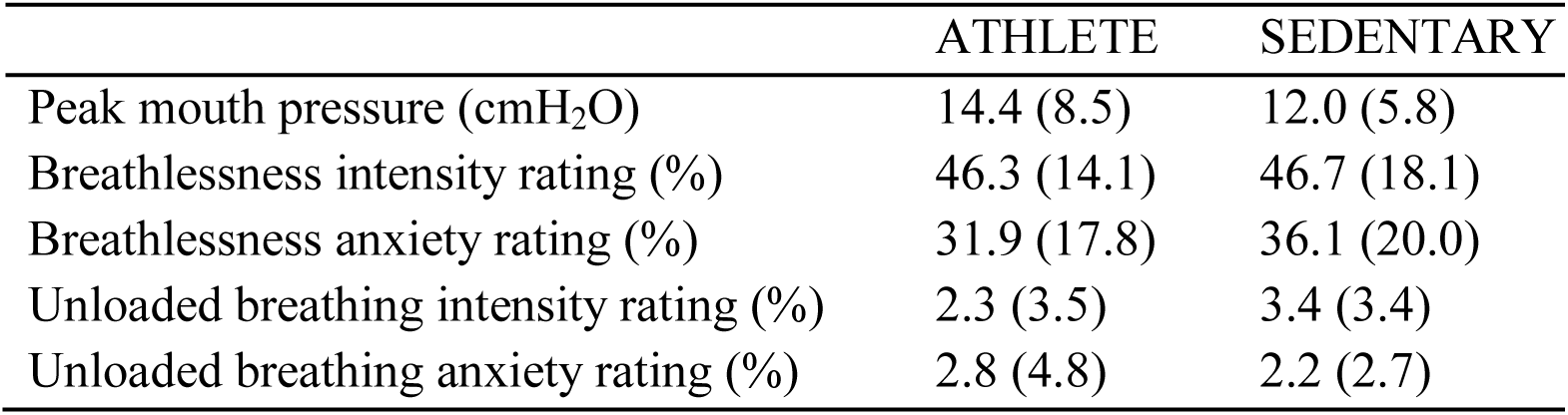
Mean (±sd) physiological and psychological variables during breathlessness for both athletes and sedentary subjects.

Two conditions were trained: 1) A shape that always predicted upcoming breathlessness (100% contingency pairing), and 2) A shape that always predicted unloaded breathing (0% contingency pairing with inspiratory resistance). The ‘certain upcoming breathlessness’ symbol was presented on the screen for 30 s, which included a varying 5-15 s anticipation period before the loading was applied. The ‘unloaded breathing’ symbol was presented for 20 s, and each condition was repeated 14 times in a randomised order. Conscious associations between cue and threat level (cue contingencies) were required and verified in all subjects by reporting (in writing) the meaning of each of the symbols both following the training session and immediately prior to the MRI scan.

Rating scores of breathing intensity were recorded after every stimulus, using a visual-analogue scale (VAS) with a sliding bar to answer the question ‘How difficult was the previous stimulus?’ where the subjects moved between ‘Not at all difficult’ (0%) and ‘Extremely difficult’ (100%). Subjects were also asked to rate how anxious each of the symbols made them feel (‘How anxious does this symbol make you feel?’) using a VAS between ‘Not at all anxious’ (0%) and ‘Extremely anxious’ (100%) immediately following the functional MRI protocol.

### Physiological measurements

We used established methods to decorrelate the effects of hypercapnia from the localised BOLD responses associated with breathing against an inspiratory resistance, using additional, matched carbon dioxide (CO_2_) boluses interspersed during rest periods in the fMRI protocols as previously described (Pattinson *et al.*, 2009*b*; Faull *et al.*, 2015; 2016*b*). In addition, a mildly hyperoxic state was achieved through a constant administration of oxygen at a rate of 0.5 L/min, to minimise fluctuations in end-tidal oxygen (P_ET_O_2_) (Table 1). Physiological measures were recorded continuously using respiratory bellows surrounding the chest, and heart rate was measured using a pulse oximeter (9500 Multigas Monitor, MR Equipment Corp., NY, USA) during the training session and MRI scan, as previously described (Faull *et al.*, 2016*b*).

### MRI scanning sequences

MRI was performed with a 7 T Siemens Magnetom scanner, with 70 mT/m gradient strength and a 32 channel Rx, single channel birdcage Tx head coil (Nova Medical).

#### BOLD scanning

A T2*-weighted, gradient echo EPI was used for functional scanning. The field of view (FOV) covered the whole brain and comprised 63 slices (sequence parameters: TE, 24 ms; TR, 3 s; flip angle, 90°; voxel size, 2 x 2 x 2 mm; field of view, 220 mm; GRAPPA factor, 3; echo spacing, 0.57 ms; slice acquisition order, descending), with 550 volumes (scan duration, 27 mins 30 s) for the task fMRI, and 190 volumes (scan duration, 9 mins 30 s) for a following resting-state acquisition (eyes open).

#### Structural scanning

A T1-weighted structural scan (MPRAGE, sequence parameters: TE, 2.96 ms; TR, 2200 ms; flip angle, 7°; voxel size, 0.7 x 0.7 x 0.7 mm; field of view, 224 mm; inversion time, 1050 ms; bandwidth; 240 Hz/Px) was acquired. This scan was used for registration of functional images.

#### Additional scanning

Fieldmap scans (sequence parameters: TE1, 4.08 ms; TE2, 5.1 ms; TR, 620 ms; flip angle, 39°; voxel size, 2 x 2 x 2 mm) of the B_0_ field were also acquired to assist distortion-correction.

### Physiological data analysis

Values for end-tidal CO_2_ (P_ET_CO_2_) were extrapolated for use as noise regressor in fMRI analysis (explained below). Respiratory waveforms, respiratory volume per unit time (RVT) and cardiac pulse oximetry triggers were included in the image denoising procedures (explained below), Values for mean and peak resistive loading, mean P_ET_CO_2_ and P_ET_O_2_, respiratory rate and RVT were calculated across each time block using custom written scripts in MATLAB (R2013a, The Mathworks, Natick, MA). Measures were averaged across each subject in each condition (unloaded breathing, anticipation and breathlessness). Peak mouth pressure was also calculated in each block and averaged in each subject for the resistive loading condition. Mean peak mouth pressure, breathlessness intensity and breathlessness anxiety ratings were then compared between the two groups using a student’s paired T-test.

### Imaging analysis

#### Preprocessing

Image processing was performed using the Oxford Centre for Functional Magnetic Resonance Imaging of the Brain Software Library (FMRIB, Oxford, UK; FSL version 5.0.8; http://www.fmrib.ox.ac.uk/fsl/). The following preprocessing methods were used prior to statistical analysis: motion correction and motion parameter recording (MCFLIRT (Jenkinson *et al.*, 2002)), removal of the non-brain structures (skull and surrounding tissue) (BET (Smith, 2002)), spatial smoothing using a full-width half-maximum Gaussian kernel of 2 mm, and high-pass temporal filtering (Gaussian-weighted least-squares straight line fitting; 120 s). B_0_ field unwarping was conducted with a combination of FUGUE and BBR (Boundary-Based-Registration; part of FEAT: FMRI Expert Analysis Tool, version 6.0 (Greve & Fischl, 2009)). Data denoising was conducted using a combination of independent components analysis (ICA) and retrospective image correction (RETROICOR) (Harvey *et al.*, 2008; Brooks *et al.*, 2013) using the externally recorded physiological measures (as previously described (Faull *et al.*, 2016*b*)), and included simultaneous regression of motion parameters.

#### Image registration

Following preprocessing, the functional scans were registered to the MNI152 (1x1x1 mm) standard space (average T1 brain image constructed from 152 normal subjects at the Montreal Neurological Institute (MNI), Montreal, QC, Canada) using a two-step process: 1) Registration of subjects’ whole-brain EPI to T1 structural image was conducted using BBR (6 DOF) with (nonlinear) fieldmap distortion-correction (Greve & Fischl, 2009), and 2) Registration of the subjects’ T1 structural scan to 1 mm standard space was performed using an affine transformation followed by nonlinear registration (FNIRT) (Andersson *et al.*, 2007).

#### Functional voxelwise and group analysis

Functional data processing was performed using FEAT (FMRI Expert Analysis Tool), part of FSL. The first-level analysis in FEAT incorporated a general linear model (Woolrich *et al.*, 2004), with the following regressors: Resistive loading periods (calculated from physiological pressure trace as onset to termination of each application of resistance); anticipation of breathlessness (calculated from onset of anticipation symbol to onset of resistance application); and unloaded breathing (onset and duration of ‘unloaded breathing’ symbol). Additional regressors to account for relief from breathlessness, periods of rating using the button box, demeaned ratings of intensity between trials, and a period of no loading following the final anticipation period (for decorrelation between anticipation and breathlessness) were also included in the analysis. A final P_ET_CO_2_ regressor was formed by linearly extrapolating between end-tidal CO_2_ peaks, and included in the general linear model to decorrelate any P_ET_CO_2_-induced changes in BOLD signal from the respiratory tasks (McKay *et al.*, 2008; Pattinson *et al.*, 2009*a*; 2009*b*; Faull *et al.*, 2015; 2016*b*). Contrasts for breathlessness (vs. baseline) and differential contrasts of anticipation of breathlessness > unloaded breathing (referred to as ‘anticipation’ or ‘anticipation of breathlessness’) were investigated at the group level.

Functional voxelwise analysis incorporated HRF modeling using three FLOBS regressors to account for any HRF differences caused by slice-timing delays, differences across the brainstem and cortex, or between individuals (Handwerker *et al.*, 2004; Devonshire *et al.*, 2012). Time-series statistical analysis was performed using FILM, with local autocorrelation correction (Woolrich *et al.*, 2001). The second and third waveforms were orthogonalised to the first to model the ‘canonical’ HRF, of which the parameter estimate was then passed up to the group analysis in a mixed-effects analysis. Group analysis was conducted using rigorous permutation testing of a General Linear Model (GLM) using FSL’s Randomize tool (Winkler *et al.*, 2014), where the GLM consisted of group mean BOLD activity for each group, and demeaned, separated breathlessness intensity and anxiety covariates for each group. Including breathlessness scores into the anticipation contrast allows us to identify preparatory brain activity that predicts the subsequent breathlessness perception when the stimulus is applied. Mean voxelwise differences between groups were calculated, as well as the interactions between group and breathlessness intensity / anxiety scores. A stringent initial cluster-forming threshold of *t* = 3.1 was used, in light of recent reports of lenient thresholding previously used in fMRI (Eklund *et al.*, 2016), and images were family-wise-error (FWE) corrected for multiple comparisons. Significance was taken at *p* < 0.05 (corrected).

#### Resting functional connectivity analysis

Following preprocessing and image registration, resting state scans from all subjects were temporally concatenated and analysed using independent component analysis (ICA) using MELODIC (Beckmann & Smith, 2004), part of FSL. ICA decomposes the data into a set of spatial maps and their associated timecourses, referred to as ‘functional networks’. Model order in the group ICA was set to 25 spatially independent components. Dual regression (Beckmann *et al.*, 2009) was then used to delineate subject-specific timecourses of these components, and their corresponding subject-specific spatial maps. Subject-specific spatial maps were again analysed non-parametrically using Randomise (part of FSL) (Winkler *et al.*, 2014) with the same GLM and significance thresholds previously applied to the functional task group analysis. Twenty components were identified as signal, and two components of interest (‘default mode’ network and ‘task positive’ network) were considered for group differences, in accordance with recent interoceptive research (Kleckner *et al.*, 2017). Therefore, *p* threshold significance was adjusted to *p* < 0.025 using Bonferroni correction for multiple comparisons.

## Results

### Physiology and psychology of breathlessness

Mean physiological values for each group for mouth pressure, P_ET_CO_2_, P_ET_O_2_, RVT, respiratory rate and RVT are presented in Table 1. Group scores for breathlessness intensity and anxiety are presented in Table 2, with no mean differences observed between groups. Previously, we have reported a difference in the accuracy between subjective breathlessness scores and changes in ventilation induced via a hypercapnic challenge (Faull *et al.*, 2016*a*) in the same subjects used as the current study. For clarity, we have reproduced the results here in Figure 4.

**Fig. 1.**
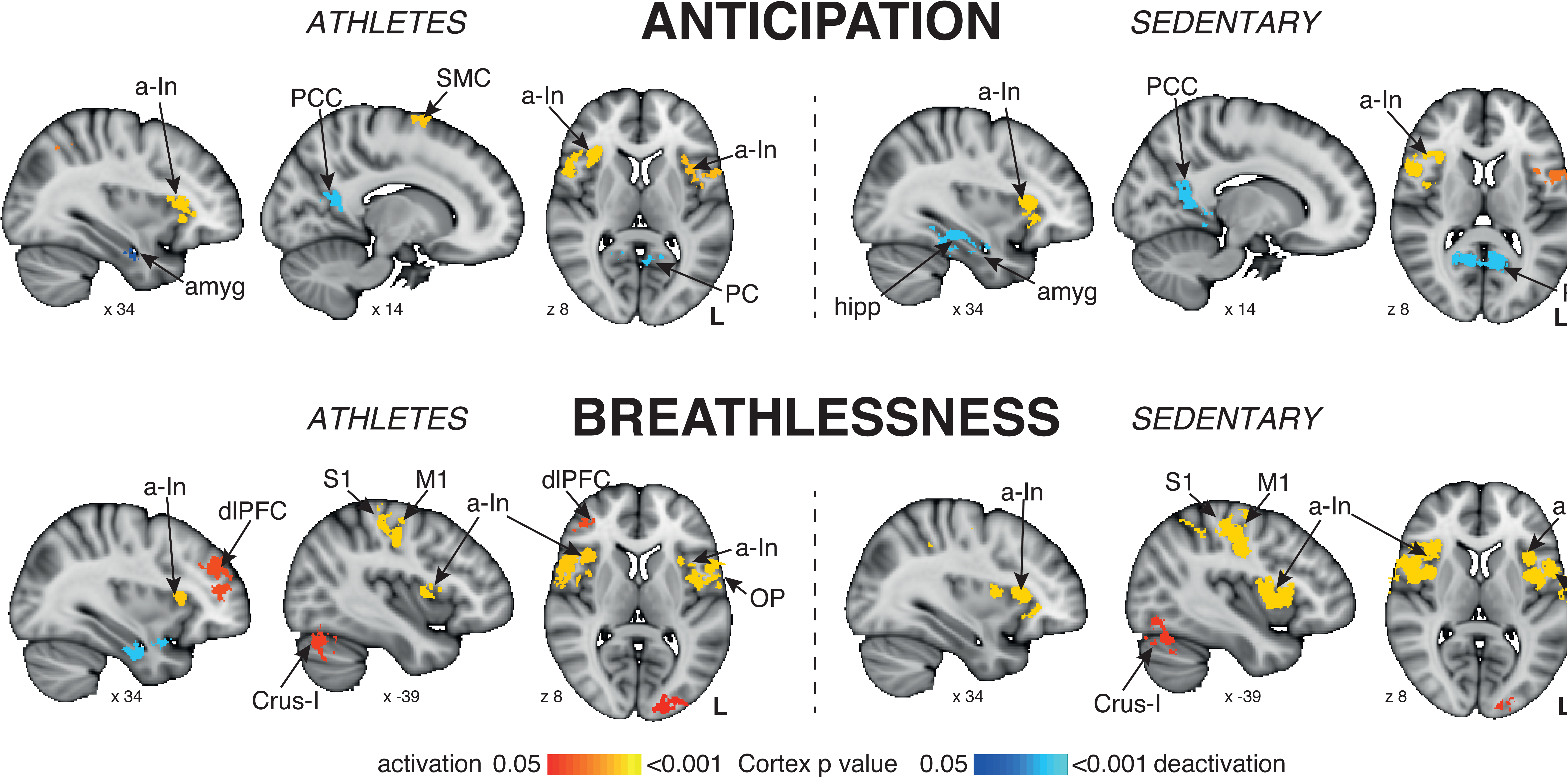
Mean BOLD activity in athletes and sedentary controls. Top: BOLD activity during conditioned anticipation of breathlessness. Bottom: BOLD activity during a breathlessness challenge, induced via inspiratory resistive loading. The images consist of a colour-rendered statistical map superimposed on a standard (MNI 1x1x1 mm) brain, and significant regions are displayed with a non-parametric cluster probability threshold of *t* < 3.1; *p* < 0.05 (corrected for multiple comparisons). Abbreviations: M1, primary motor cortex; SMC, supplementary motor cortex; dACC, dorsal anterior cingulate cortex; PCC, posterior cingulate cortex; dlPFC, dorsolateral prefrontal cortex; a-In, anterior insula; OP, operculum; amyg, amygdala; hipp, hippocampus; Crus-I, cerebellar lobe; activation, increase in BOLD signal; deactivation, decrease in BOLD signal.

**Fig. 2.**
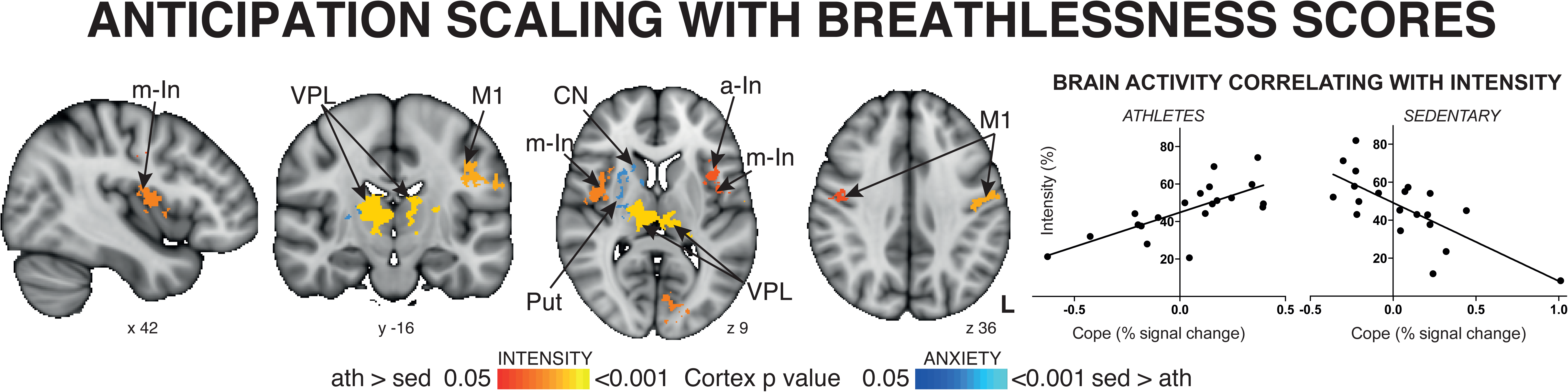
Interaction between groups and breathlessness scores. Left: BOLD activity during conditioned anticipation of breathlessness. Red-yellow = BOLD activity correlating with intensity scores in athletes > sedentary subjects; blue-light blue = BOLD activity correlating with anxiety scores in sedentary > athletic subjects. Right: Percentage BOLD signal change within the (red-yellow) intensity-correlated imaging mask against intensity scores, demonstrating a positive, linear correlation in athletes and a negative relationship in sedentary subjects. The images consist of a colour-rendered statistical map superimposed on a standard (MNI 1x1x1 mm) brain, and significant regions are displayed with a non-parametric cluster probability threshold of *t* < 3.1; *p* < 0.05 (corrected for multiple comparisons). Abbreviations: M1, primary motor cortex; a-In, anterior insula; m-In, middle insula; hipp, hippocampus; put, putamen; CN, caudate nucleus; VPL, ventral posterolateral thalamic nucleus.

**Fig. 3.**
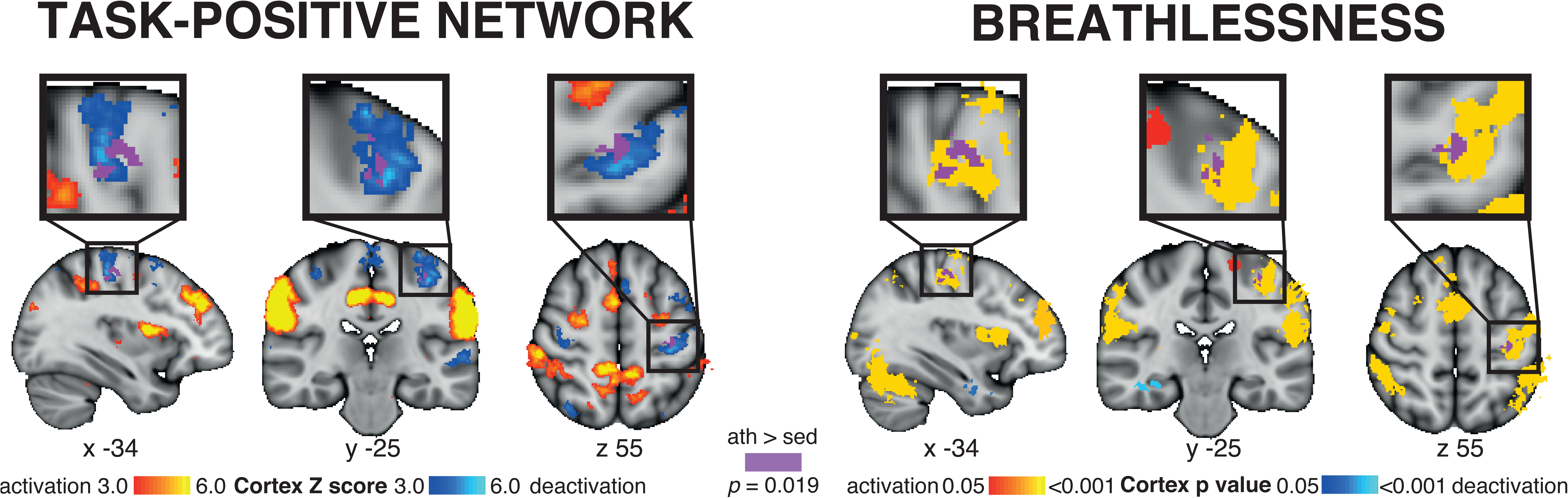
Differences in resting functional connectivity between athletes and sedentary subjects. Increased functional connectivity (purple) observed in athletes between an area of primary motor corticex that is active during breathlessness (right) and a cingulo-opercular task-positive network (left) identified at rest. The images consist of a colour-rendered statistical map superimposed on a standard (MNI 1x1x1 mm) brain, and significant regions are displayed with a non-parametric cluster probability threshold of *t* <3.1; *p* < 0.05 (corrected for multiple comparisons).

**Fig. 4.**
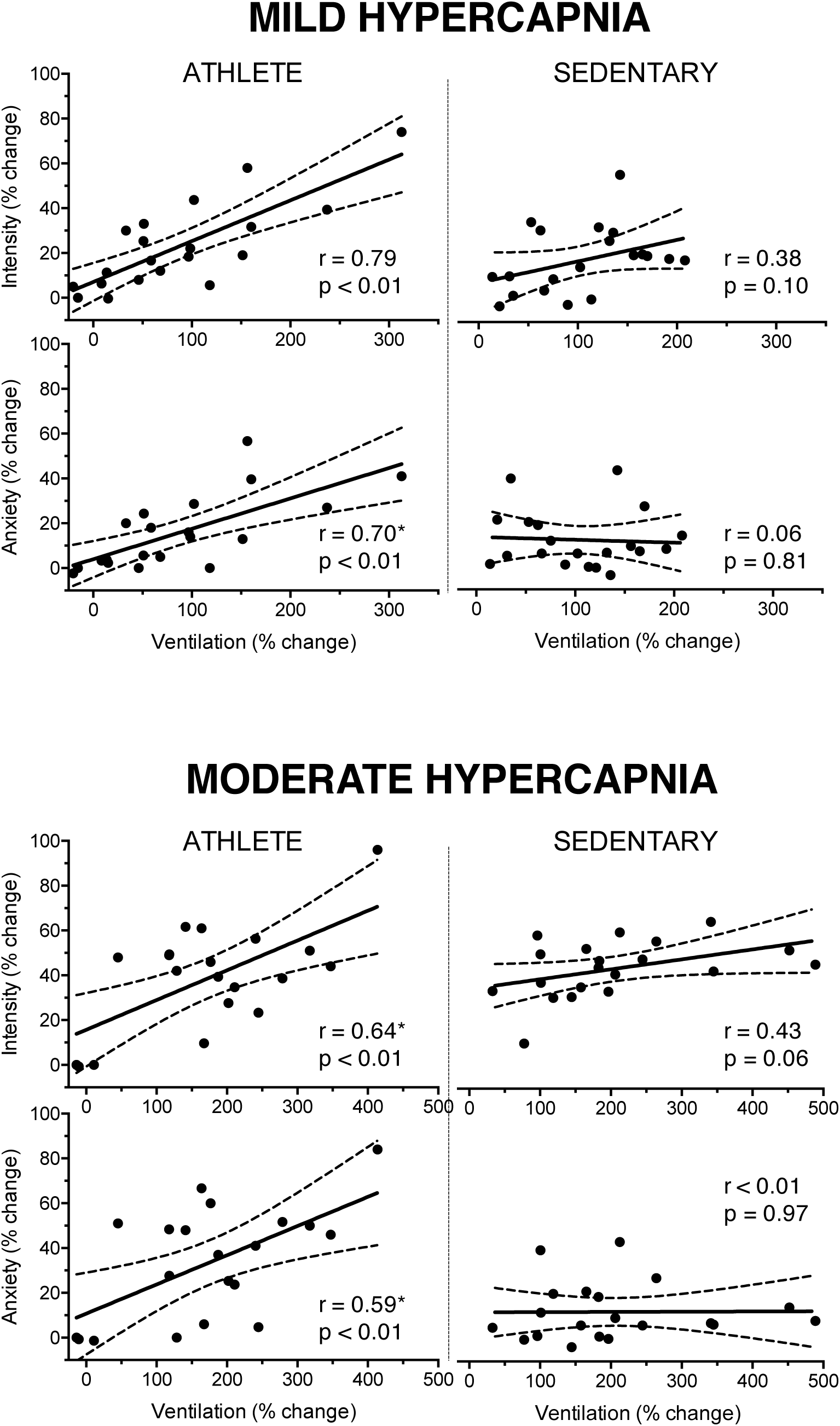
Previously reported differences in ventilatory perceptions between athletes and sedentary subjects. *Significantly different slope from sedentary subjects. Subject-specific change in breathlessness anxiety and intensity scores plotted against percentage change in ventilation from baseline, induced by both mild (top) and moderate (bottom) hypercapnia (mild hypercapnia: aim of 0.8%; and moderate hypercapnia: aim of 1.5% increase in end-tidal carbon dioxide, end-tidal oxygen was maintained constant). Athletes are plotted in the left column, and sedentary subjects in the right column. During both mild and moderate hypercapnia, the athlete group showed a positive linear correlation between change in ventilation and change in breathing anxiety that was significantly different from sedentary subjects (slope difference: mild p = 0.018; moderate p = 0.011). Athletes also demonstrated significant positive correlations for breathlessness intensity against change in ventilation, where the slope was significantly different to sedentary subjects in moderate (p = 0.047) but not mild (p = 0.177) hypercapnia. 95% Confidence intervals are shown. Figure is recreated from previously published data (Faull *et*, 2016*a*) under the Creative Commons license.

### Task fMRI analysis

#### Mean group differences

Mean activity during anticipation of breathlessness in each group is presented in Figure 1. In sedentary subjects, significantly increased BOLD activity was observed in the right anterior insula, operculum and bilateral primary motor cortex, and decreased BOLD activity in bilateral posterior cingulate cortex, precuneus, lateral occipital cortex, hippocampus, parahippocampal gyrus and amygdala. In athletes, increased BOLD activity was observed in bilateral anterior insula, operculum and primary motor cortex, and right supplementary motor cortex, and decreased BOLD activity in bilateral precuneus, hippocampus, parahippocampal gyrus and amygdala. No statistically significant voxelwise differences were observed between group mean activities during anticipation of breathlessness (differentially contrasted against unloaded breathing).

Mean activity during breathlessness in each group is presented in Figure 1. In sedentary subjects, significantly increased BOLD activity was observed in the bilateral anterior and middle insula, operculum, primary sensory and motor cortices, supplementary motor cortex, supramarginal gyrus and cerebellar VI, and decreased BOLD activity in bilateral precuneus. In athletes, significantly increased BOLD activity was observed in the right dorsolateral prefrontal cortex, bilateral anterior and middle insula, operculum, primary sensory and motor cortices, supplementary motor cortex, left visual cortex and cerebellar Crus-I, and decreased BOLD activity in right amygdala, hippocampus and superior temporal gyrus. No statistically significant voxelwise differences were observed between group mean activities during breathlessness.

#### Subjective breathlessness scores

The brain activity that correlated with breathlessness scores of intensity and anxiety was compared between groups, to identify any interaction effects (group x subjective score). Interaction effects establish that the difference between groups varies as a function of the covariate (subjective scores). Athletes demonstrated widespread brain activity positively correlating with (predicted) intensity scores during anticipation of breathlessness (Figure 2), whilst those same areas had a negative correlation in sedentary subjects (interaction). This included activity in the bilateral ventral posterolateral nucleus of the thalamus, middle insula, and primary motor and sensory cortices, as well as left anterior insula. In contrast, a small amount of activity in the right putamen and caudate nucleus correlated with anxiety in sedentary subjects, but not in athletes during anticipation. No significant interactions between groups were present for either intensity of anxiety during breathlessness perception.

### Resting state network connectivity

Of the 25 resting state ‘networks’ identified in the group ICA analysis, 20 components were identified to represent relevant signal (19 cortical, 1 cerebellar) while the remaining 5 were labeled as noise (see supplementary material for a summary the 20 resting networks). Two networks of interest were identified for group comparison analyses (as determined by Kleckner et al., 2017): 1) The network most representative of the typical ‘default mode’, and 2) A network containing components of previously identified visual and dorsal attention networks (Vossel *et al.*, 2014), which was notably most similar to the breathing task contrasts (‘task-positive’ network) (Figure 3). When network connectivity was compared between athletes and controls, athletes were found to have significantly greater (*p* = 0.019) connectivity of the task-positive network to an area of primary motor cortex active during resistive loading (Figure 3).

## Discussion

### Main findings

We have identified a cohesive anticipatory brain network that predicts upcoming subjective ratings of breathlessness in athletes. Comparatively, this brain activity was reversed (i.e. negatively correlated with upcoming breathlessness ratings) in sedentary controls. Furthermore, at rest, athletes demonstrated greater connectivity between an area of breathing-related primary sensorimotor cortex, and a cingulo-opercular attention network that is strikingly similar to that recently identified to be involved in allostatic-interoceptive processing (Kleckner *et al.*, 2017). This network may therefore be integral within attention and processing of sensory signals related to breathing. Increased connectivity between sensorimotor cortex and this brain network may underlie the observed differences in anticipatory processing of respiratory signals, and the improved ventilatory perceptive accuracy found in these endurance athletes.

### Breathlessness processing in athletes

Endurance athletes have repeated episodes of elevated ventilation and perceptions of breathlessness as part of their training. In previously published results (Faull *et al.*, 2016*a*), we have demonstrated improved psychophysical matching between changes in chemostimulated hyperventilation and subjective breathlessness perceptions in these athletes compared to matched sedentary subjects (Figure 4). Therefore, whether by nature or nurture, these individuals appear to have improved ventilatory perception accuracy. The reduced correlation between changes in ventilation and perceptions of breathlessness demonstrated in sedentary subjects implies a worsened ability to process respiratory sensations, which may be a risk factor for symptom discordance in disease (Van den Bergh *et al.*, 2017).

In accordance with behavioural findings, here we have observed differences in the brain processing of breathing perceptions in athletes. Specifically, a coherent network of brain activity corresponding to breathlessness intensity scores was observed during anticipation in athletes, which was reversed (negatively correlated) with subjective scores in sedentary subjects. This network incorporates key areas involved in sensorimotor control and interoception, such as the thalamus, insula and primary sensorimotor cortices (Feldman & Friston, 2010; Simmons *et al.*, 2012; Feldman Barrett & Simmons, 2015; Van den Bergh *et al.*, 2017). The opposing relationship between brain activity and subjective scores in athletes and sedentary subjects indicates a fundamental difference in preparatory, anticipatory brain activity directed towards subjective perceptions between these groups, which occurs without any difference in overall group mean brain activity. Conversely, sedentary subjects demonstrated activity corresponding to anxiety scores in the ventral striatum (caudate nucleus and putamen) during anticipation of breathlessness. The striatum has been previously linked with cardiovascular responses resulting from social threat (Wager *et al.*, 2009), and may represent heightened threat responses in sedentary subjects.

Interestingly, the intensity-related differences in brain activity were observed during the anticipation period that preceded the actual perception of breathlessness. It is possible that repeated increases in ventilation and breathlessness during training helps athletes improve the accuracy of their breathing expectations for an upcoming stimulus, such as expecting to run up a hill. Recent theories of symptom perception have proposed a comprehensive, Bayesian model (Feldman Barrett & Simmons, 2015; Van den Bergh *et al.*, 2017), which includes a set of perceptual expectations or ‘priors’. These expectations are combined with sensory information from the periphery, for the brain to probabilistically produce the most likely resulting perception. Furthermore, factors such as attention (Merikle & Joordens, 1997; Phelps *et al.*, 2006; Ling & Carrasco, 2006) and interoceptive ability (Gray *et al.*, 2007; Critchley *et al.*, 2013; Mallorqui-Bague *et al.*, 2016; Garfinkel *et al.*, 2016*b*) are thought to influence this system, either by altering the prior expectations or incoming sensory information.

While previous research had identified reduced anterior insula activity during loaded breathing in endurance athletes (Paulus *et al.*, 2012), we have not reproduced these findings when employing more stringent fMRI statistics. Nevertheless, the proposal by Paulus and colleagues (Paulus *et al.*, 2012) that athletes demonstrate more efficient minimization of the body prediction error remains a very plausible possibility. Here, instead, we have observed functional perception-related differences during anticipation of loaded breathing in endurance athletes. Therefore, repeated exercise training in athletes may develop breathlessness expectations (or priors) and better direct attention towards breathing sensations, improving the robustness of the perceptual system to accurately infer the intensity of breathlessness.

### Differences in functional connectivity within the athletic brain

Understanding differences in underlying communication between functional brain regions may inform us as to why differences in functional activity, such as observed in these athletes during anticipation of breathlessness, may arise. The temporal synchronicity of seemingly spontaneous fluctuations in brain activity across spatially distinct regions can inform us of how ‘functionally connected’ these disparate regions may be, and is thought to be related to the temporal coherence of neuronal activity in anatomically distinct areas (Gerstein & Perkel, 1969; Van Den Heuvel & Pol, 2010).

It is now well established that the brain can be functionally parsed into resting state ‘networks’, where distinct brain regions are consistently shown to exhibit temporally similar patterns of brain activity (Smith *et al.*, 2009; Miller *et al.*, 2016). While properties of these resting state networks have been linked to lifestyle, demographic and psychometric factors (Smith *et al.*, 2015; Miller *et al.*, 2016), here we have found connectivity differences between athletes and sedentary subjects for a cingulo-opercular network. This network displays a very similar spatial distribution to the pattern of activity observed during the breathlessness tasks (‘task-positive’) (Figure 3), as well as the allostatic-interoceptive network recently identified by Kleckner and colleagues (Kleckner *et al.*, 2017), and to previously reported networks of ventral and dorsal attention (Fox *et al.*, 2005; 2006). Here, we have demonstrated greater functional connectivity in athletes between an area of primary sensory and motor cortices that has consistently been identified as active during tasks such as breath holds (Pattinson *et al.*, 2009*b*; Faull *et al.*, 2015) and inspiratory resistances (Faull *et al.*, 2016*b*; Faull & Pattinson, 2017; Hayen *et al.*, 2017). Therefore, it is possible that this greater connectivity in athletes between an interoceptive attention network and primary sensorimotor cortex contributes to the processing of incoming and outgoing respiratory information, and thus may also be related to more accurate ventilatory perceptions.

Whilst this cross-sectional study is unable to determine whether endurance exercise training *induces* these differences in brain function and connectivity, or whether these individuals are biased towards training for endurance sports, this work provides intriguing preliminary insight that the brain may undergo adaptation in conjunction with the periphery, to more accurately process perceptions of bodily sensations such as breathlessness.

### Neuroimaging statistical considerations

Extensive efforts were made within the analysis of this dataset to ensure only the most robust and reliable results were reported. Firstly, physiological noise and potential motion artifacts need to be specifically addressed when using breathing-related tasks, and these can be further compounded at higher field strengths (Brooks et al., 2013). Here we employed rigorous noise correction procedures, combining retrospective image correction of physiological parameters (heart rate, ventilation and end-tidal carbon dioxide) with both extended motion parameter regression and independent component analysis de-noising (Faull et al., 2016b; Hayen et al., 2017). Secondly, recent work has revealed the potential leniency of previous fMRI statistical methodologies and thresholds (Eklund et al., 2016). In this manuscript, we have utilized minimal (2mm) spatial smoothing to maintain accurate localization of brain activity, and employed non-parametric, permutation testing with a robust cluster threshold of 3.1 (Eklund et al., 2016), to represent only the most reliable statistical results. Whilst these approaches forsake much of our previously-reported activity within these breathing-related tasks (Faull & Pattinson, 2017), we can have greater confidence in our reported differences between brain and behavior in athletes and sedentary subjects.

### Potential clinical implications of altering breathlessness processing

As discussed, prior expectations of breathlessness are now considered to be a major contributor to symptom perception (Hayen *et al.*, 2013; Faull *et al.*, 2017; Van den Bergh *et al.*, 2017; Geuter *et al.*, 2017; Herigstad *et al.*, 2017). Altering the accuracy of breathlessness perception using exercise training may be of interest when treating individuals with habitual symptomology, such as those with chronic obstructive pulmonary disease (COPD) or asthma. Recent research has shown exercise training to reduce breathlessness intensity and anxiety in patients with COPD, with corresponding changes in the brain’s processing of breathlessness-related words (Herigstad *et al.*, 2016; 2017). It has been proposed that exercise exposure alters breathlessness expectations and priors in these patients, modifying symptom perception when it has become discordant with physiology in chronic disease (Parshall *et al.*, 2012; Herigstad *et al.*, 2017). It is also possible that exercise helps improve the processing of respiratory signals for more accurate ventilatory interoception in these patients, allowing breathlessness perception to better match respiratory distress. Future work investigating the link between exercise, ventilation and breathlessness perception may yield another treatment avenue (via targeted exercises) to improve patient quality of life in the face of chronic breathlessness.

## Conclusions

In this study, we have demonstrated altered anticipatory brain processing of breathlessness intensity in athletes compared to sedentary subjects. This altered functional brain activity may be underpinned by increased functional connectivity between an interoceptive network related to breathlessness, and sensorimotor cortex that is active during ventilatory tasks. These differences in brain activity and connectivity may also relate to improvements in ventilatory perception previously reported between these subject groups (Faull *et al.*, 2016*a*), and open the door to investigating exercise as a tool to manipulate brain processing of debilitating breathing symptoms, such as breathlessness in clinical populations.

## Acknowledgements

This research was supported by the JABBS Foundation. This research was further supported by the National Institute for Health Research, Oxford Biomedical Research Centre based at Oxford University Hospitals NHS Trust and University of Oxford. Olivia K Faull was supported by the Commonwealth Scholarship Commission.

## References

Andersson JL, Jenkinson M & Smith S (2007). Non-linear registration, aka Spatial normalisation FMRIB technical report TR07JA2. FMRIB Analysis Group of the University of Oxford.

Beckmann CF & Smith SM (2004). Probabilistic Independent Component Analysis for Functional Magnetic Resonance Imaging. IEEE Transactions on Medical Imaging 23, 137–137.

Beckmann CF, Mackay CE, Filippini N & Smith SM (2009). Group comparison of resting-state FMRI data using multi-subject ICA and dual regression. NeuroImage 47, S148.

Bogaerts K, Notebaert K, Van Diest I, Devriese S, De Peuter S & Van den Bergh O (2005). Accuracy of respiratory symptom perception in different affective contexts. J Psychosom Res 58, 537–537.

Borg E, Borg G, Larsson K, Letzter M & Sundblad BM (2010). An index for breathlessness and leg fatigue. Scand J Med Sci Sports 20, 644–644.

Brooks JCW, Faull OK, Pattinson KTS & Jenkinson M (2013). Physiological noise in brainstem FMRI. Front Hum Neurosci 7, 623–623.

Carrieri-Kohlman V, Gormley JM, Douglas MK, Paul SM & Stulbarg MS (1996). Exercise training decreases dyspnea and the distress and anxiety associated with it: monitoring alone may be as effective as coaching. Chest Journal 110, 1526–1526.

Carrieri-Kohlman V, Gormley JM, Eiser S, Demir-Deviren S, Nguyen H, Paul SM & Stulbarg MS (2001). Dyspnea and the affective response during exercise training in obstructive pulmonary disease. Nursing Research 50, 136–136.

Critchley HD, Eccles J & Garfinkel SN (2013). Interaction between cognition, emotion, and the autonomic nervous system. In Autonomic Nervous System, Handbook of Clinical Neurology, pp. 59–59. Elsevier.

Davenport PW & Vovk A (2009). Cortical and subcortical central neural pathways in respiratory sensations. Respiratory physiology & neurobiology 167, 72–72.

Devonshire IM, Papadakis NG, Port M, Berwick J, Kennerley AJ, Mayhew JEW & Overton PG (2012). Neurovascular coupling is brain region-dependent. NeuroImage 59, 1997–1997.

Eklund A, Nichols TE & Knutsson H (2016). Cluster failure: Why fMRI inferences for spatial extent have inflated false-positive rate. PNAS 113, 7900–7900.

El-Manshawi A, Killian KJ, Summers E & Jones NL (1986). Breathlessness during exercise with and without resistive loading. Journal of Applied Physiology 61, 896–896.

Faull O, Hayen A & Pattinson K (2017). Breathlessness and the body: Neuroimaging clues for the inferential leap. Cortex; DOI: 10.1101/117408.

Faull OK & Pattinson KT (2017). The cortical connectivity of the periaqueductal gray and the conditioned response to the threat of breathlessness. Elife 6, 95.

Faull OK, Cox PJ & Pattinson KTS (2016a). Psychophysical Differences in Ventilatory Awareness and Breathlessness between Athletes and Sedentary Individuals. Front Physiol 7, 195–195.

Faull OK, Jenkinson M, Clare S & Pattinson KTS (2015). Functional subdivision of the human periaqueductal grey in respiratory control using 7 tesla fMRI. NeuroImage 113, 356–356.

Faull OK, Jenkinson M, Ezra M & Pattinson KTS (2016b). Conditioned respiratory threat in the subdivisions of the human periaqueductal gray. Elife; DOI: 10.7554/elife.12047.

Feldman Barrett LF & Simmons WK (2015). Interoceptive predictions in the brain. Nat Rev Neurosci 16, 419–419.

Feldman H & Friston KJ (2010). Attention, Uncertainty, and Free-Energy. Front Hum Neurosci 4, 1–1.

Fox MD, Corbetta M, Snyder AZ, Vincent JL & Raichle ME (2006). Spontaneous neuronal activity distinguishes human dorsal and ventral attention systems (vol 103, pg 10046, 2006). Proceedings of the National Academy of Sciences 103, 13560–13560.

Fox MD, Snyder AZ, Vincent JL, Corbetta M, Van Essen DC & Raichle ME (2005). The human brain is intrinsically organized into dynamic, anticorrelated functional networks. Proceedings of the National Academy of Sciences 102, 9673–9673.

Garfinkel SN, Manassei MF, Hamilton-Fletcher G, In den Bosch Y, Critchley HD & Engels M (2016a). Interoceptive dimensions across cardiac and respiratory axes. Philos Trans R Soc Lond, B, Biol Sci 371, 20160014–20160014.

Garfinkel SN, Tiley C, O’Keeffe S, Harrison NA, Seth AK & Critchley HD (2016b).Discrepancies between dimensions of interoception in autism: Implications for emotion and anxiety. Biological Psychology 114, 117–117.

Gerstein GL & Perkel DH (1969). Simultaneously recorded trains of action potentials: analysis and functional interpretation. Science 164, 828–828.

Geuter S, Boll S, Eippert F & Büchel C (2017). Functional dissociation of stimulus intensity encoding and predictive coding of pain in the insula. Elife 6, e24770.

Gray MA, Harrison NA, Wiens S & Critchley HD (2007). Modulation of Emotional Appraisal by False Physiological Feedback during fMRI. PLoS ONE 2, e546.

Greve DN & Fischl B (2009). Accurate and robust brain image alignment using boundary-based registration. NeuroImage 48, 63–63.

Handwerker DA, Ollinger JM & D’Esposito M (2004). Variation of BOLD hemodynamic responses across subjects and brain regions and their effects on statistical analyses. NeuroImage 21, 1639–1639.

Harvey AK, Pattinson KTS, Brooks JCW, Mayhew SD, Jenkinson M & Wise RG (2008).Brainstem functional magnetic resonance imaging: Disentangling signal from physiological noise. Journal of Magnetic Resonance Imaging 28, 1337–1337.

Hayen A, Herigstad M & Pattinson KTS (2013). Understanding dyspnea as a complex individual experience. Maturitas 76, 45–45.

Hayen A, Wanigasekera V, Faull OK, Campbell SF, Garry PS, Raby SJM, Robertson J, Webster R, Wise RG, Herigstad M & Pattinson KTS (2017). Opioid suppression of conditioned anticipatory brain responses to breathlessness. NeuroImage 150, 383–383.

Herigstad M, Faull O, Hayen A, Evans E, Hardinge M, Wiech K & Pattinson KTS (2017). Treating breathlessness via the brain: Mechanisms underpinning improvements in breathlessness with pulmonary rehabilitation. European Respiratory Journal; DOI: 10.1101/117390.

Herigstad M, Hayen A, Reinecke A & Pattinson KTS (2016). Development of a dyspnoea word cue set for studies of emotional processing in COPD. Respiratory physiology & neurobiology 223, 37–37.

Holloszy JO & Coyle EF (1984). Adaptations of Skeletal-Muscle to Endurance Exercise and Their Metabolic Consequences. Journal of Applied Physiology 56, 831–831.

Jenkinson M, Bannister P, Brady M & Smith S (2002). Improved Optimization for the Robust and Accurate Linear Registration and Motion Correction of Brain Images. NeuroImage 17, 825–825.

Jones AM & Carter H (2000). The Effect of Endurance Training on Parameters of Aerobic Fitness. Sports Medicine 29, 373–373.

Kaufman MP & Forster HV (1996). Reflexes Controlling Circulatory, Ventilatory and Airway Responses to Exercise. John Wiley & Sons, Inc., Hoboken, NJ, USA.

Kleckner IR, Zhang J, Touroutoglou A, Chanes L, Xia C, Simmons WK, Quigley KS, Dickerson BC & Feldman Barrett L (2017). Evidence for a large-scale brain system supporting allostasis and interoception in humans. Nat hum behav 1, 0069–0069.

Lang PJ, Wangelin BC, Bradley MM, Versace F, Davenport PW & Costa VD (2011). Threat of suffocation and defensive reflex activation. Psychophysiol 48, 393–393.

Lansing RW, Im B, Thwing JI, Legedza A & Banzett RB (2000). The perception of respiratory work and effort can be independent of the perception of air hunger. Am J Respir Crit Care Med 162, 1690–1690.

Ling S & Carrasco M (2006). When sustained attention impairs perception. Nat Neurosci 9, 1243–1243.

Mallorqui-Bague N, Bulbena A, Pailhez G, Garfinkel SN & Critchley HD (2016). Mind-Body Interactions in Anxiety and Somatic Symptoms. Harvard Review of Psychiatry 24, 53–53.

McKay LC, Adams L, Frackowiak RSJ & Corfield DR (2008). A bilateral cortico-bulbar network associated with breath holding in humans, determined by functional magnetic resonance imaging. NeuroImage 40, 1824–1824.

Merikle PM & Joordens S (1997). Parallels between Perception without Attention and Perception without Awareness. Consciousness and cognition 6, 219–219.

Miller KL et al. (2016). Multimodal population brain imaging in the UK Biobank prospective epidemiological study. Nat Neurosci 19, 1523–1536.

Parshall MB, Schwartzstein RM, Adams L, Banzett RB, Manning HL, Bourbeau J, Calverley PM, Gift AG, Harver A, Lareau SC, Mahler DA, Meek PM & O’Donnell DE (2012). An Official American Thoracic Society Statement: Update on the Mechanisms, Assessment, and Management of Dyspnea. Am J Respir Crit Care Med 185, 435–435.

Pattinson K, Mitsis GD, Harvey AK & Jbabdi S (2009a). Determination of the human brainstem respiratory control network and its cortical connections in vivo using functional and structural imaging. NeuroImage 44, 295–295.

Pattinson KTS, Governo RJ, MacIntosh BJ, Russell EC, Corfield DR, Tracey I & Wise RG (2009b). Opioids Depress Cortical Centers Responsible for the Volitional Control of Respiration. Journal of Neuroscience 29, 8177–8177.

Paulus MP, Flagan T, Simmons AN, Gillis K, Kotturi S, Thom N, Johnson DC, Van Orden KF, Davenport PW & Swain JL (2012). Subjecting Elite Athletes to Inspiratory Breathing Load Reveals Behavioral and Neural Signatures of Optimal Performers in Extreme Environments ed. Lucia A. PLoS ONE 7, e29394.

Pavlov IP (1927). Conditioned Reflexes. Phelps EA, Ling S & Carrasco M (2006). Emotion facilitates perception and potentiates the perceptual benefits of attention. Psychol Sci 17, 292–292.

Porro CA, Baraldi P, Pagnoni G, Serafini M, Facchin P, Maieron M & Nichelli P (2002). Does anticipation of pain affect cortical nociceptive systems? The Journal of Neuroscience 22, 3206–3206.

Price DD, Milling LS, Kirsch I, Duff A, Montgomery GH & Nicholls SS (1999). An analysis of factors that contribute to the magnitude of placebo analgesia in an experimental paradigm. Pain 83, 147–147.

Simmons WK, Avery JA, Barcalow JC, Bodurka J, Drevets WC & Bellgowan P (2012). Keeping the body in mind: Insula functional organization and functional connectivity integrate interoceptive, exteroceptive, and emotional awareness. Human brain mapping 34, 2944–2958.

Smith SM (2002). Fast robust automated brain extraction. Human brain mapping 17, 143–155.

Smith SM, Fox PT, Miller KL, Glahn DC, Fox PM, Mackay CE, Filippini N, Watkins KE, Toro R, Laird AR & Beckmann CF (2009). Correspondence of the brain’s functional architecture during activation and rest. 1–1.

Smith SM, Nichols TE, Vidaurre D, Winkler AM, Behrens TEJ, Glasser MF, Ugurbil K, Barch DM, Van Essen DC & Miller KL (2015). A positive-negative mode of population covariation links brain connectivity, demographics and behavior. Nature Med 18, 1565–1565.

Spinhoven P, vanPeskiOosterbaan AS, VanderDoes A, Willems L & Sterk PJ (1997).Association of anxiety with perception of histamine induced bronchoconstriction in patients with asthma. Thorax 52, 149–149.

Takano N, Inaishi S & Zhang Y (1997). Individual differences in breathlessness during exercise, as related to ventilatory chemosensitivities in humans. The Journal of Physiology 499, 843–848.

Tang J & Gibson S (2005). A Psychophysical Evaluation of the Relationship Between Trait Anxiety, Pain Perception, and Induced State Anxiety. The Journal of Pain 6, 612–612.

Van den Bergh O, Witthöft M, Petersen S & Brown RJ (2017). Symptoms and the body: Taking the inferential leap. Neuroscience & Biobehavioral Reviews 74, 185–185.

Van Den Heuvel MP & Pol HEH (2010). Exploring the brain network: a review on resting-state fMRI functional connectivity. European Neuropsychopharmacology 20, 519–519.

Vossel S, Geng JJ & Fink GR (2014). Dorsal and Ventral Attention Systems: Distinct Neural Circuits but Collaborative Roles. Neuroscientist 20, 150–150.

Wager TD, Rilling JK, Smith EE, Sokolik A, Casey KL, Davidson RJ, Kosslyn SM, Rose RM & Cohen JD (2004). Placebo-Induced Changes in fMRI in the Anticipation and Experience of Pain. Science 303, 1162–1162.

Wager TD, Waugh CE, Lindquist M, Noll DC, Fredrickson BL & Taylor SF (2009). Brain mediators of cardiovascular responses to social threat. NeuroImage 47, 821–821.

Waldrop TG, Eldridge FL, Iwamoto GA & Mitchell JH (2010). Central Neural Control of Respiration and Circulation During Exercise, 2nd edn. John Wiley & Sons, Inc., Hoboken, NJ, USA.

Winkler AM, Ridgway GR, Webster MA, Smith SM & Nichols TE (2014). Permutation inference for the general linear model. NeuroImage 92, 381–381.

Woolrich MW, Behrens TEJ, Beckmann CF, Jenkinson M & Smith SM (2004). Multilevel linear modelling for FMRI group analysis using Bayesian inference. NeuroImage 21, 1732–1732.

Woolrich MW, Ripley BD, Brady M & Smith SM (2001). Temporal Autocorrelation in Univariate Linear Modeling of FMRI Data. NeuroImage 14, 1370–1370.

